# ADAGE signature analysis: differential expression analysis with data-defined gene sets

**DOI:** 10.1101/156620

**Authors:** Jie Tan, Matthew Huyck, Dongbo Hu, René A. Zelaya, Deborah A. Hogan, Casey S. Greene

## Abstract

**Background:** Gene set enrichment analysis and overrepresentation analyses are commonly used methods to determine the biological processes affected by a differential expression experiment. This approach requires biologically relevant gene sets, which are currently curated manually, limiting their availability and accuracy in many organisms without extensively curated resources. New feature learning approaches can now be paired with existing data collections to directly extract functional gene sets from big data.

**Results:** Here we introduce a method to identify perturbed processes. In contrast with methods that use curated gene sets, this approach uses signatures extracted from public expression data. We first extract expression signatures from public data using ADAGE, a neural network-based feature extraction approach. We next identify signatures that are differentially active under a given treatment. Our results demonstrate that these signatures represent biological processes that are perturbed by the experiment. Because these signatures are directly learned from data without supervision, they can identify uncurated or novel biological processes. We implemented ADAGE signature analysis for the bacterial pathogen *Pseudomonas aeruginosa*. For the convenience of different user groups, we implemented both an R package (**ADAGEpath**) and a web server (http://**adage.greenelab.com**) to run these analyses. Both are open-source to allow easy expansion to other organisms or signature generation methods. We applied ADAGE signature analysis to an example dataset in which wild-type and *Δanr* mutant cells were grown as biofilms on the Cystic Fibrosis genotype bronchial epithelial cells. We mapped active signatures in the dataset to KEGG pathways and compared with pathways identified using GSEA. The two approaches generally return consistent results; however, ADAGE signature analysis also identified a signature that revealed the molecularly supported link between the MexT regulon and Anr.

**Conclusions:** We designed ADAGE signature analysis to perform gene set analysis using data-defined functional gene signatures. This approach addresses an important gap for biologists studying non-traditional model organisms and those without extensive curated resources available. We built both an R package and web server to provide ADAGE signature analysis to the community.

## Background

High-throughput genome-scale measurements are now widely used because they can provide a global view of a biological system. Typical experiments involve a control and some sort of treatment, and the typical output is a list of genes with expression levels that were significantly altered. In addition to examining genes of interest individually, researchers often summarize results with gene set overrepresentation analysis (also called pathway analysis) to infer the biological basis of the gene list. These analyses aim to link groups of perturbed genes by their biological themes and help researchers understand the effect of an experiment on biological pathways.

Two primary components comprise gene set analyses: a testing algorithm and pre-defined sets of biologically themed genes. While the first part has been extensively explored [1–3], the second part has not drawn much attention because the creation and maintenance of gene sets remains a largely manual process requiring substantial curator effort. Currently, gene sets are primarily contributed by consortia of curators, such as the GO consortium [4,5]. Manual annotation ensures the quality of the gene sets but is slow, can be tedious, and leads to gene sets with certain biases [6]. Furthermore, while a small set of primary model organisms has received substantial curator effort, other organisms remain sparsely annotated. Accurately transferring annotations across organisms using computational prediction algorithms remains challenging, particularly for biological processes [7]. Due to the limited availability and sparse coverage of gene sets, the potential of gene set analysis remains limited for most non-traditional model organisms.

In contrast to the paucity of carefully curated gene sets specific to these non-traditional models, the amount of genome-wide gene expression data has grown rapidly, especially for microbes which have a relatively small transcriptome and are inexpensive to assay [8]. For single-cell organisms, a complete compendium of public data ideally captures expression under numerous conditions. We may expect these compendia to extensively characterize many of the organism’s transcriptional regulatory processes and to be well-suited targets for the extraction of pathway-like signatures.

We previously developed ADAGE, an algorithm that extracts meaningful gene sets from genome-wide gene expression compendia [9]. ADAGE models are unsupervised neural network models of large publicly available gene expression compendia. Specifically, ADAGE models are denoising autoencoder neural networks [10,11]. ADAGE first encodes gene expression data into neural network nodes and then decodes those nodes to reconstruct the original expression levels. The denoising term in the name comes from the fact that random noise is added to the input while encoding individual samples. This results in a model that is able to reconstruct expression levels without noise and has been shown to make models more robust [11]. These neural network nodes describe features of the input data that capture essential patterns, i.e. those robust to noise. Our analysis of the genes that most influence each node previously revealed that they form gene sets that resemble human-annotated biological processes and pathways, which often exhibit consistent coexpression in large gene expression compendia [9,12]. We have termed such gene sets ADAGE signatures. We developed eADAGE, which summarizes multiple ADAGE models into an ensemble model, to more robustly capture pathways and found that it covered significantly more biological pathways more precisely [12]. In addition to signatures that match curated pathways, eADAGE also extracts signatures that group genes that match known but uncurated pathways and others that may represent undiscovered biological processes.

To fully leverage signatures built by eADAGE or other robust feature construction approaches, we introduce an ADAGE signature analysis pipeline. ADAGE signature analysis aims to identify one or more signatures that respond to an experimental treatment. As with gene set analyses, these signatures represent biological processes that may be perturbed by the treatment. The approach is similar to traditional gene set analysis but replaces human-annotated gene sets with ADAGE-learned signatures. ADAGE signature analysis complements pathway-style analysis in any organism by providing an unsupervised perspective, and is usable for non-traditional model organisms or other organisms for which curated pathways are unavailable. Here we demonstrate ADAGE signature analysis in the bacterial pathogen *Pseudomonas aeruginosa*. We chose *P. aeruginosa* as our model organism because it has sufficient public gene expression data to construct a model and a dedicated research community. Though sparsely curated in the recent past, its pathway annotations have been growing rapidly due to a community annotation initiative [13]. This allows us to validate the biological relevance of gene signatures learned by ADAGE, while also demonstrating its ability to identify as yet unannotated biological processes. To facilitate the use of ADAGE signature analysis, we developed both an R package for users with bioinformatics background and an easy-to-use web server intended for use by bench biologists.

## Methods

### ADAGE signature analysis workflow

ADAGE signature analysis has three major steps: data preparation, active signature detection, and signature interpretation (Figure 1). In addition to the input dataset to be analyzed, ADAGE signature analysis also requires an ADAGE model and the gene expression compendium of an organism from which the model was built. The *P. aeruginosa* compendium and (e)ADAGE models can be built following instructions in [9,12].

**Figure 1:**
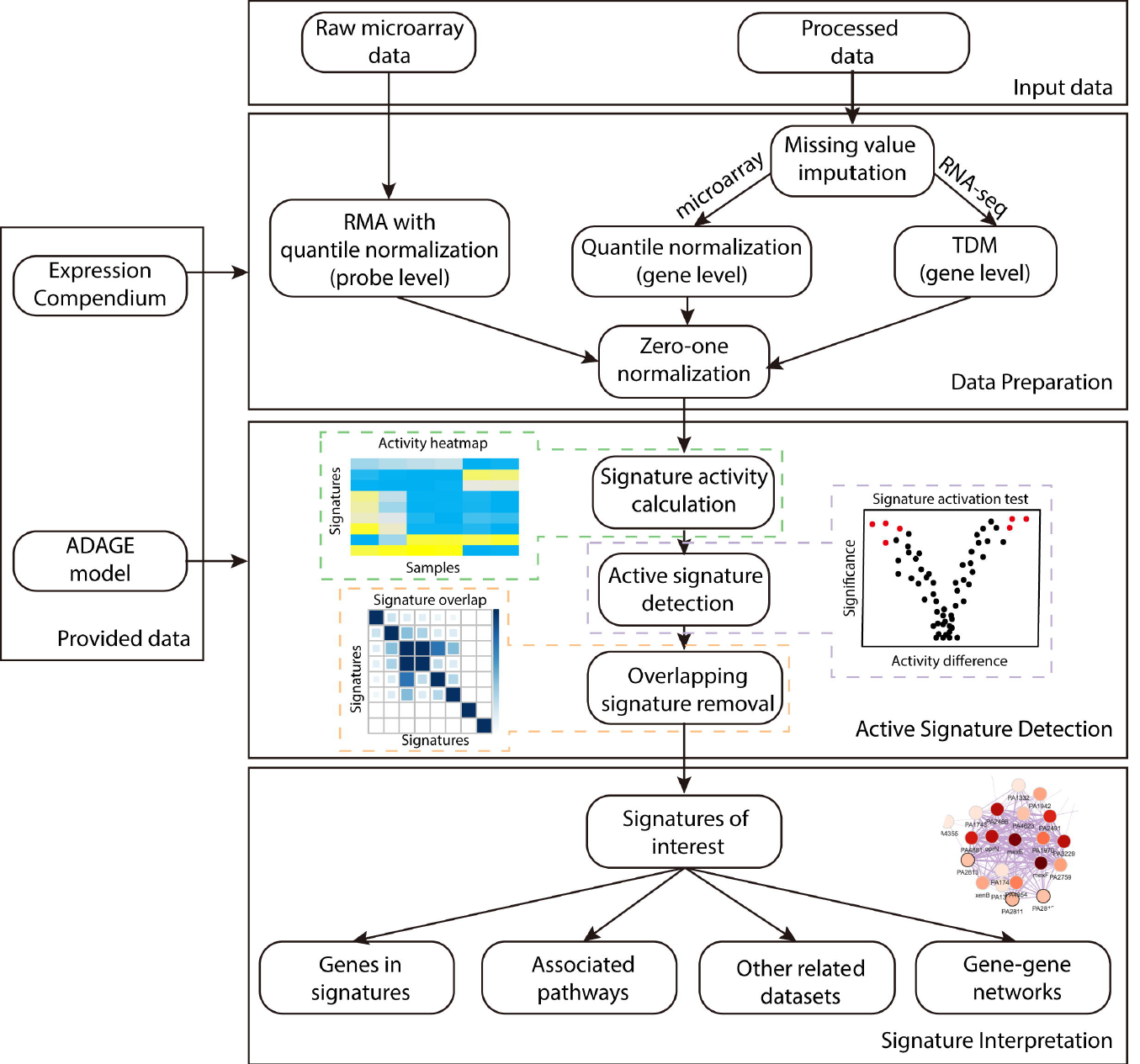
ADAGE signature analysis workflow. An input dataset is first processed to be compatible with a provided ADAGE model trained on the provided expression compendium. Raw microarray measurements are transformed by RMA, while processed data first have missing values imputed and then are normalized to the compendium. The second step is detecting active signatures in the dataset. Signature activity for each sample in the dataset is computed. Statistical test on the signature activity is used to identify signatures respond to an experimental treatment. Redundant signatures are further removed. The last step is interpreting the biological meaning of active signatures through evaluating genes in the signatures, pathways associated with the signatures, datasets related to the signatures, and a gene-gene network built upon the signatures.

#### Data preparation

Gene expression data must be normalized to be compatible with the ADAGE model. Raw microarray data measured on the same chip platform as the compendium are analyzed with Robust Multi-array Average (RMA) [14]. RMA includes background correction, quantile normalization, and probe summarization; however, quantile normalization is performed against the quantile distribution of the compendium so that the resulting expression values are comparable with the compendium. Other types of gene expression data must first be processed into gene-level expression values. Gene identifiers used in the input data are mapped to the gene identifiers used in the compendium. We next impute the expression of missing genes using k-nearest neighbors - the neighbors are computed based on similarity in the compendium. For processed microarray data, we apply quantile normalization using the compendium’s quantiles. For RNA-seq data, expression values are normalized to the compendium via TDM [15]. The last step in data preparation for all types of input data is a zero-one linear transformation using the compendium as reference. Measurements outside the range observed in the compendium are set to zero or one according to whether they are below or above the range. After processing, the dataset is ready for ADAGE signature analysis.

#### Active signature detection

The concept of ADAGE signature was first introduced in [12]. To recap, in an ADAGE model, genes connect to nodes via weights and this vector of weights characterizes each node (Figure 2). The distribution of the weight vector centers near zero and is close to normal, with a small number of genes contributing high weights. We group genes with weights more extreme than 2.5 standard deviations from the mean on the positive side and the negative side of the weight distribution separately. These two gene sets form the positive signature and negative signature of a node and are simply named as “NodeXXpos” and “NodeXXneg”. The positive and negative signatures of the same node are not always related. As opposed to a simple set of genes, the signature is a gene set with a weight value for each gene that indicates the gene’s importance to the set. For simplicity and because the positive and negative sides of the distribution are already separated, we ignore the signs of weight values.

**Figure 2:**
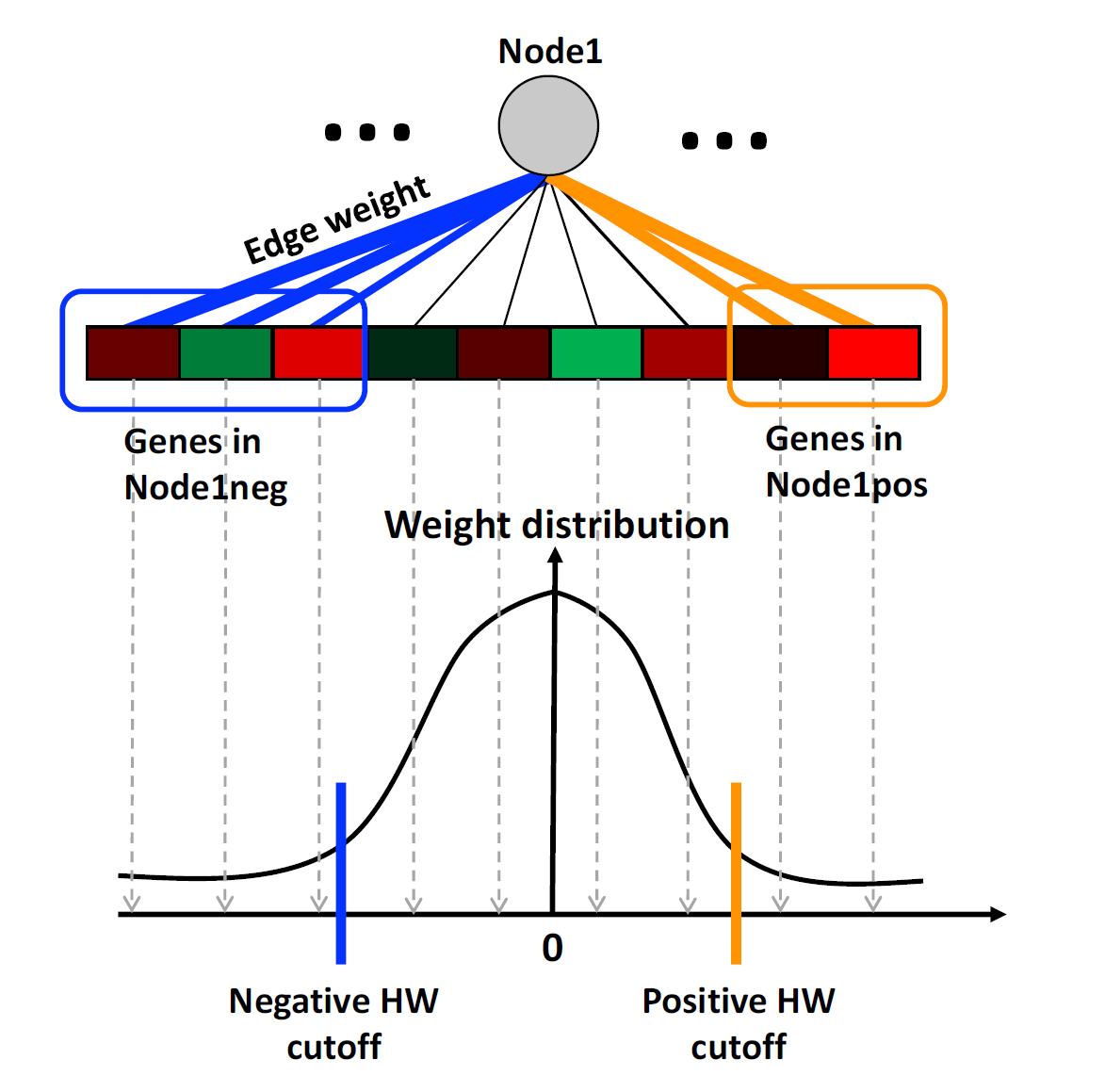
ADAGE model and gene signatures. In an ADAGE model, every gene is linked to every node through an edge. The edge weight is fixed after model training and its magnitude is reflected by edge thickness. The distribution of gene weights to a node is centered at zero and close to normal. Genes giving weights higher than the positive high-weight (HW) cutoff together form the positive gene signature for that node (genes in the orange circle). Similarly, genes giving weights lower than the negative HW cutoff together form negative gene signature for that node (genes in the blue circle).

After data preparation, we calculate each signature’s activity for every sample in the input dataset. The signature’s activity reflects how active that signature is in each sample and is defined as the average expression values of signature genes weighted by genes’ absolute weights in the signature. This results in a matrix of activity values, where each row is a signature and each column is a sample, which can be shown in an activity heatmap (Figure 1). To detect signatures associated with an experimental treatment, we apply statistical tests to signature activities. The most appropriate statistical test depends on the experimental design. After applying the selected test, we can identify a list of the signatures with the greatest changes in activity in response to a particular experimental treatment.

We have observed that multiple signatures can share many genes. Signature overlap may be due in part to the random noise added during training; however, we cannot rule out the possibility that the subtle differences between two overlapping signatures are biologically meaningful. Therefore, we must carefully handle signature overlap to remove redundant signatures but retain those relevant to a specific dataset. To accomplish this, we calculate marginal activities of every combination of two signatures. The marginal activity for a signature pair is the activity of one signature after removing genes that it overlaps with the second signature. After removal, we test whether the marginal activities still respond significantly to the treatment. We ignore signatures when their activity is no longer significant after removing the effect of another signature, as long as the other signature is not also removed through this process. In a special case where a group of signatures all become non-significant after removing each other, we keep the one that is most significantly altered. This process results in a final list of signatures affected by an experimental treatment.

#### Signature interpretation

It is important to note that a benefit of ADAGE signature analysis, as opposed to attempting to interpret the entire set of signatures, is that investigators only need to examine signatures that are affected by their experiment. Signatures are gene sets formed based on the expression patterns in a gene expression compendium. They are not annotated to a specific biological process, but we have used several strategies to help understand the biological meaning behind a signature. The most intuitive way is to examine its gene composition. If some genes have been previously characterized and share a biological theme, they may suggest the biological process represented by the signature. Users can also link existing curated biological knowledge such as KEGG [16] pathways and GO terms [17] to signatures through an enrichment test, though such annotations are not always available. Even when they exist, annotations are not expected to be comprehensive for non-traditional model organisms. Finally, users can probe a signature by analyzing the compendium and extracting experiments in which the signature has its largest activity ranges. We expect these to be experimental conditions in which the biological processes represented by the signature are perturbed. Taken together or individually, these steps allow researchers to quickly interpret signatures of interest without curating a complete collection of pathways.

To visualize which genes are in signatures and how they are related across the overall model, we construct a gene-gene network using genes in selected signatures of interest. The network is built upon gene-gene relationships extracted from an ADAGE model. In the network, two genes are linked by an edge if the correlation between their weight vectors, i.e. how strongly connected they are to each node, is higher than a tunable cutoff. Depending on how they are linked with each other, genes can form modules in the network. These modules highlight functional units of genes in differentially active signatures. The network can be interactively explored and its use is facilitated by overlaid information, such as gene descriptions, differential expression in the experiment being analyzed, and annotations for each gene from GO and KEGG where available.

### User interface

There are two ways for users to access ADAGE signature analysis. We provide an R package, intended for computationally inclined users and a web server intended for those without familiarity with the R programming language. The R package and the web server are both preloaded with a *Pseudomonas aeruginosa* gene expression compendium containing microarray samples measured on the Pae_G1a Affymetrix *Pseudomonas aeruginosa* array that were available on the ArrayExpress database [18] before July 31 2015, a previously published eADAGE model built on this compendium [12], and *P.a.* gene information retrieved from NCBI’s ftp site. Both are open source and licensed freely, so that investigators can add their own machine learning models and additional organisms. We also plan to expand both resources to include additional non-traditional model organisms.

#### Web server

We developed a web server that implements the most central components of ADAGE signature analysis. The web server is designed with a clean separation between a backend API and a JavaScript application frontend. This allows programmatic access to the server if desired. The backend is written in Python using the Django framework. The frontend uses AngularJS to provide a responsive user experience, with Vega and D3 used to provide interactive visualizations. Both the backend and frontend are available under the permissive 3-clause BSD open source license. Advanced users can initialize their own instance of the ADAGE web server, load models of their choosing, and supply this interface to users. Our public instance of the ADAGE web server is hosted on Amazon Web Services. Here we describe the main features provided by the web server. Users first need to choose a machine learning model on the homepage and all the following analyses are model specific. Then users can explore assays and experiments and perform signature analysis (Analyze), explore genes’ similarities in the model through a gene-gene network (GeneNetwork), explore signatures in the model (Signature), and obtain annotations for the underlying sample compendium (Download).

### Analyze

The Analyze feature guides users to perform the entire ADAGE signature analysis pipeline (Figure 3). To begin the analysis, users first search experiments or samples with a keyword. Next users identify experiments or samples of interest and add them to the analysis basket. Clicking the analysis basket takes users to the Sample page, where sample information is listed. A heatmap is plotted showing signature activities in all samples and can be clustered by sample or signature. To compare two groups of samples, users assign samples to either a treatment (yellow) or a control (purple) group. Then a differential activation test between the two groups is performed and its result is presented in a volcano plot showing difference in mean activity in the x-axis and significance p-value in the y-axis. Signatures with positive activity differences are active in the comparison group. Users choose signatures that are highly differentially active and examine them either in the Signature page or the GeneNetwork page described below.

**Figure 3:**
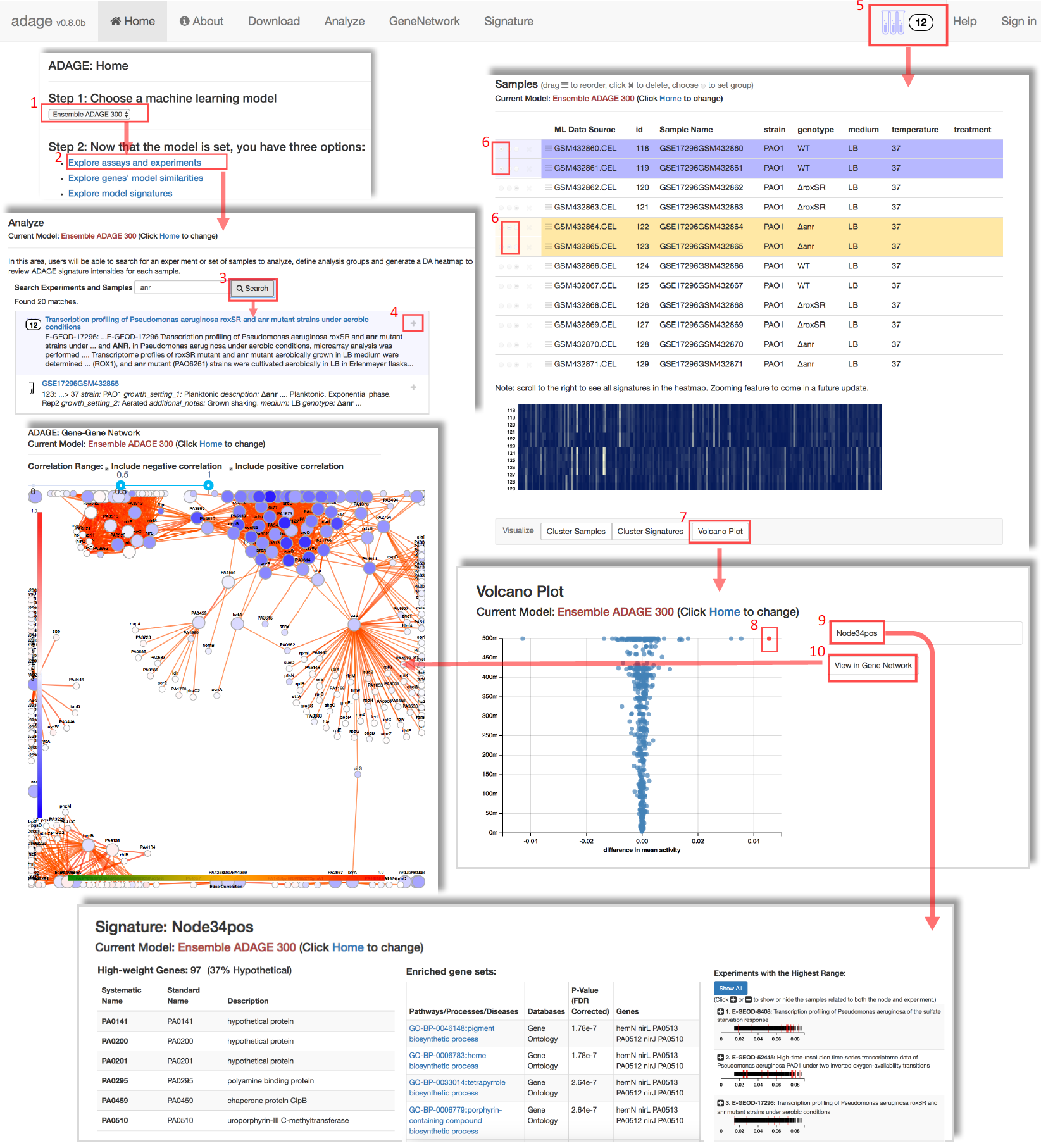
ADAGE web server interface and analysis workflow. The ADAGE web server interface has six tabs on the top. The signature analysis pipeline starts with choosing a machine learning model on the homepage (step1) and choosing to explore assays and experiments (step2). In the Analyze page, users can search datasets or samples using keywords (step3). After clicking the “+” button beside each dataset or sample, the dataset with all samples in it or an individual sample is added into the analysis basket (step4). After users add samples, clicking the basket brings users to the Sample page (step5). The Sample page provides experimental information about each sample and a signature activity heatmap. Users can define a two-group comparison by selecting samples and assigning them to either treatment (yellow) group or control (purple) group (step6). The signatures that are differentially active between two groups of samples can be examined in a volcano plot (step7). Next users select signatures in the volcano plot (step8) and further inspect them in the Signature page (step9). The Signature page provides information about gene composition, gene set association, and related datasets of a signature. Lastly, users can visualize their interested signatures in a gene-gene network (step10). Users can also directly examine genes in the gene-gene network through the GeneNetwork tab and examine a signature through the Signature tab.

### GeneNetwork

The GeneNetwork feature allows users to investigate a gene or a group of genes in an ADAGE-derived gene-gene network. Users input genes of interest from the organism associated with that machine learning model. A network including input genes and their connecting genes drawn from the machine learning model will be shown. The default view presents genes connected by an absolute edge correlation higher than 0.5. Users can adjust the correlation range to examine more strongly or weakly connected genes. Another way to obtain genes of interest is from a two-group comparison through the Analysis feature as described above. If the gene-gene network results from this process, then the color of each gene in the network reflects gene expression fold change. Otherwise, the node color is grey. Gene information such as names and function description are provided when clicking on a node. Signatures shared by two connecting genes are shown when clicking on an edge.

### Signature

The Signature feature helps users to interpret the biological meaning of an ADAGE signature. After choosing a machine learning model, users can directly examine a signature in the model. The Signature page lists each gene in a signature and its known functions. It also presents a table of GO and KEGG pathways associated with the signature. To help identify other public datasets that have the signature active, it also shows experiments with the highest activity range for that signature.

### Download

The Download feature allows users to download sample annotations. These annotations are manually curated experimental information for each sample in the training compendium [12].

#### R package

We built an R package called *ADAGEpath* to perform ADAGE signature analysis. It is written exclusively in R [19] using the devtools package [20] and is available on github (https://github.com/greenelab/ADAGEpath) under the BSD-3-clause license. Here we describe the main functions in the package.

The function *load_dataset()* loads and processes an input dataset from local machine or directly from the ArrayExpress database if an accession number is provided. The recommended input format is a set of raw CEL files, which can be directly processed from the probe level with the help of the *affy* [21], *affyio* [22], and *preprocessCore* [23] packages from Bioconductor. When CEL files are not available or the input dataset is measured by RNAseq, *load_dataset()* also accepts processed expression values at the gene level. Missing values in processed data are imputed with the *impute* Bioconductor package. RNAseq measurements are further transformed using the *TDM* package [24]. Since ADAGE uses expression values between 0 and 1, input expression values are linearly transformed to be between 0 and 1 using the function *zeroone_norm()*.

The function *calculate_activity()* calculates each signature’s activity for each sample in the dataset. To identify signatures with differential activities, we suggest using the Bioconductor package *limma* [25], particularly when sample size is small. *Limma* builds linear models to test differential activation and estimates variance more robustly than the t-test. To facilitate the most frequently used two-group comparison, we wrap a limma-based two-group differential activation test into the function *build_limma()*. The test significance and the activity difference between two groups can be visualized using *plot_volcano()*. The function *get_active_signatures()* supports three ways to prioritize most activated signatures: filtering by significance, sorting by absolute activity difference, and optimizing significance and activity difference simultaneously through choosing signatures lying on the top Pareto fronts. In multi-objective optimization, if there is no other solution that outperforms a specific solution over all objectives, that solution is said to be Pareto optimal. All such solutions make up the Pareto front. *Limma* also supports many types of experimental designs, such as factorial design and time-course experiments. We provide examples of analyzing a time-course experiment and a factorial-design experiment using *limma* in the package vignettes. Users can also apply other statistical tests to identify activated signatures if desired. The function *plot_activity_heatmap()* generates a heatmap showing how signature activity changes across samples.

To remove redundant signatures, users can calculate the marginal activity for each permutation of two signatures using *calculate_marginal_activity()*. When the comparison is between two groups of samples, the function *plot_marginal_activation()* helps visualize whether a signature is still strongly active after the impact of another signature has been removed and the function *remove_redundant_signatures()* returns non-redundant active signatures for a dataset. Users can check how signatures overlap with each other in their gene compositions using the function *plot_signature_overlap()*.

To get a detailed view of a signature or a group of signatures, users can retrieve their constituent genes using *annotate_genes_in_signatures()*. Users can also download existing GO terms and KEGG pathways [16] from the TRIBE web server [26] using *fetch_geneset()* and associate signatures to known GO and KEGG pathways using *annotate_signatures_with_genesets()*. The TRIBE web server also allows users to build and share their own custom gene sets, so the connection to this resource enables custom gene set analysis as well. Lastly, users can visualize signatures via gene-gene networks using the function *visualize_gene_network()*. The network is built with the help of the R package *igraph* [27] and made interactive with the package *visNetwork* [28].

ADAGEpath is built upon many existing R packages. In addition to the packages mentioned above, ADAGEpath uses functions from *gplots* [29], *corrplot* [30], *leaflet* [31], and *plotly* [32] for plotting; *httr* [33] and *jsonlite* [34] for data querying; *readr* [35], *data.table* [36], *tibble* [37], *dplyr* [38], *magrittr* [39], *R.utils* [40], and *reshape2* [41].

#### A comparison between two user interfaces

The web server and R package target users with different backgrounds and needs. For bench scientists, the web server is straightforward to use and does not require familiarity with any programming language. For bioinformaticians, the R package provides more flexibility. It allows programmatic access and integration with other analysis pipelines. Powered by rich statistical resources in R, the R package tests the differential activation using the more robust linear model (provided by the limma R package). The web server currently supports the two-sample t-test for differential activation. Therefore, it’s normal to get slightly different results from the two platforms. To avoid security and privacy issues with data storage and management, the web server currently supports only public datasets. The overlapping signature removal function is not available in the web server at this time. Users who wish to automatically filter overlapping signatures or analyze datasets that are not publicly available should use the R package.

## Results

Here we demonstrate ADAGE signature analysis on an example dataset (GSE67006). This dataset contains wild-type and *Δanr* mutant grown as biofilms on the Cystic Fibrosis genotype bronchial epithelial cells (CFBE) in order to model cystic fibrosis airways infections. Anr is a transcriptional regulator responsible for the aerobic to anaerobic transition [42]. We performed ADAGE signature analysis to identify biological processes that were affected by Anr on CFBE cells. The script that reproduces the following analysis is available on Github (https://github.com/greenelab/SignatureAnalysis-CaseStudy).

We first ran a two-group limma test between wild-type samples and mutant samples to detect signatures with significantly different activities. We visualized test results as a volcano plot with activity difference on the x-axis and test significance on the y-axis (Figure 4A). Many signatures passed the 0.05 significance cutoff. To focus on the most differentially active signatures, we considered both activity differences and statistical significance by selecting signatures that were on the top 10 layers of Pareto fronts. This resulted in 36 signatures and their activities in each sample were visualized with a heatmap in which yellow indicated high activity and blue indicated low activity (Figure 4B).

**Figure 4:**
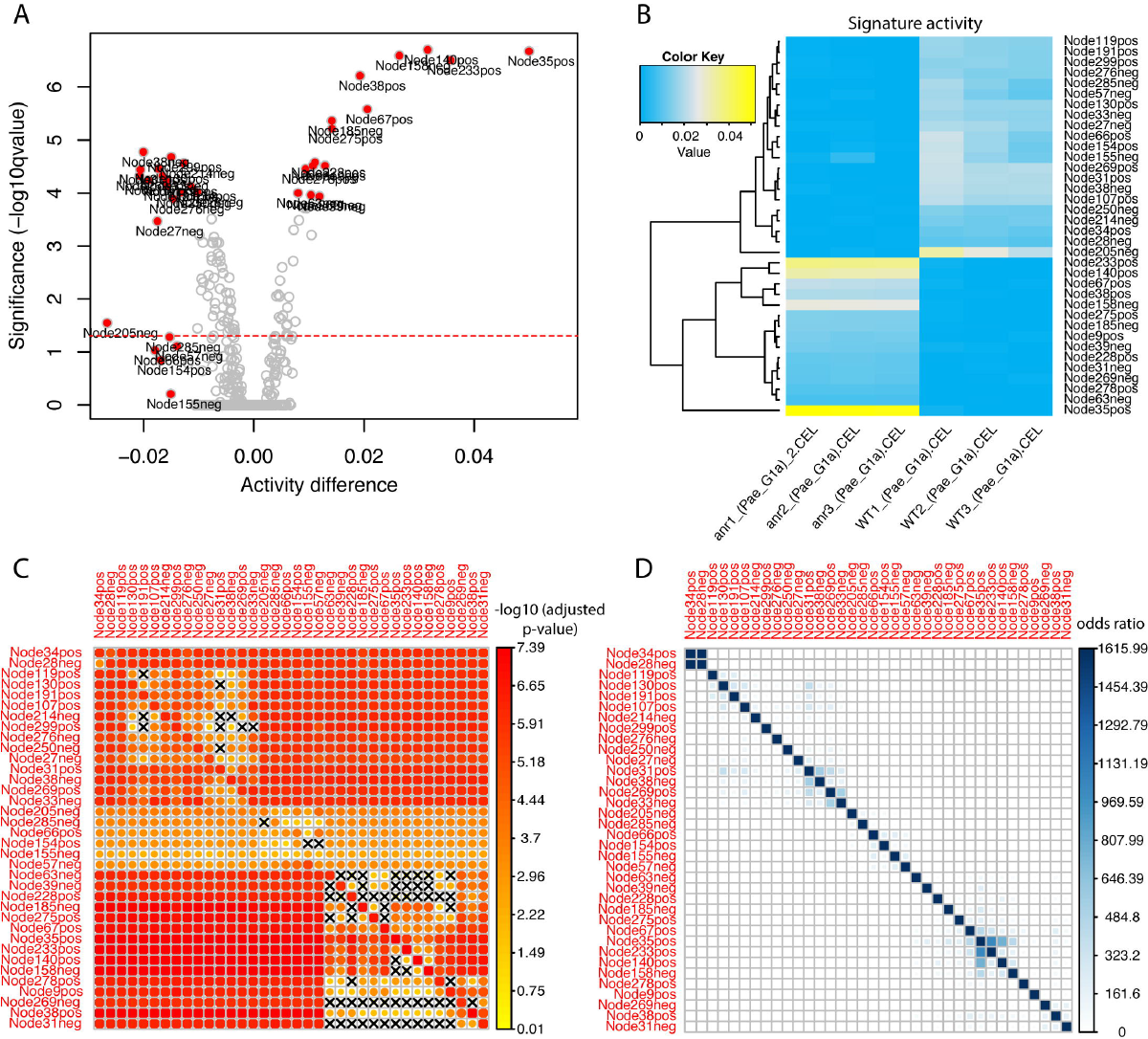
ADAGE signature analysis for the example dataset. A: A volcano plot showing both the activity difference and the statistical test significance in the activation test between *Δanr* mutant and wild-type. Signatures that lie on the top 10 layers of Pareto fronts are labeled and highlighted in red. The red dotted line indicates the significance of –log10(0.05). B: A heatmap showing the absolute activity difference of signatures detected in panel A. C: A non-symmetric heatmap reflecting the marginal activation significance (adjusted p value in –log10 scale) between every signature pair in panel A. For two signatures, one in the row and the other in the column, the heatmap color depicts how strong the signature in the row is differentially activated after the genes it shares with the signature in the column have been depleted. A cross sign indicates a non-significant activation (adjusted p-value > 0.05). The diagonal shows the activation significance of a signature without overlapping gene removal. D: A symmetric heatmap reflecting signature similarity between every signature pair in panel A. The heat color represents the odds ratio that two signatures overlap in their gene contents. Because comparing a signature with itself would result in an odds ratio of infinity, values on the diagonal are replaced by the highest odds ratio among all signature pairs.

Because signatures could overlap in their gene compositions, we evaluated the effects of ADAGE parameters on overlap. We observed that signatures tend to have more overlapping genes when the model is trained with higher corruption level (the amount of noise added) (Figure S1A, B). For each overlap, we cannot rule out the possibility that subtle differences are biologically meaningful. Supporting the potential biological distinctions between similar signatures, models trained with more noise also cover more biological pathways within a reasonable corruption range (0% to 25% corruption) (Figure S1C). For this reason, we employed a data-driven process to remove overlapping signatures before performing detailed investigation of specific signatures.

We calculated the marginal activity for each combination of signature pairs and tested its significance in differentiating the wild-type and deletion strains. Figure 4C shows the test significance (adjusted p-values in the -log10 scale) when the signature in each column was removed from the signature in the row and the diagonal shows the significance of each signature. A cross sign indicates non-significant p-values. We define a signature to be redundant if it becomes non-significant after removing the effect of another signature. Following these rules, we dropped the following signatures: Node119pos, Node214neg, Node299pos, Node130pos, Node250neg, Node154pos, Node63neg, Node39neg, Node228pos, Node158neg, Node140pos, Node269neg, Node31neg, Node185neg, Node275pos, Node278pos, and Node285neg. Interestingly, Node34pos and Node28neg shared many genes (Figure 4D), but they each contained additional genes and both remained significant in the marginal activation test. This result highlighted the importance of viewing signature overlap in the context of a dataset. At this stage, there were 19 differentially active signatures remaining. These were visualized together in a gene-gene network (edge correlation cutoff = 0.5) (Figure 5A); the edges between genes in this network revealed sets of genes that have transcriptional relationships as detected in the ADAGE model. Network modules, discrete clusters within the network, can reveal regulons. For example, the sets of genes involved in denitrification (Figure 5C).

**Figure 5:**
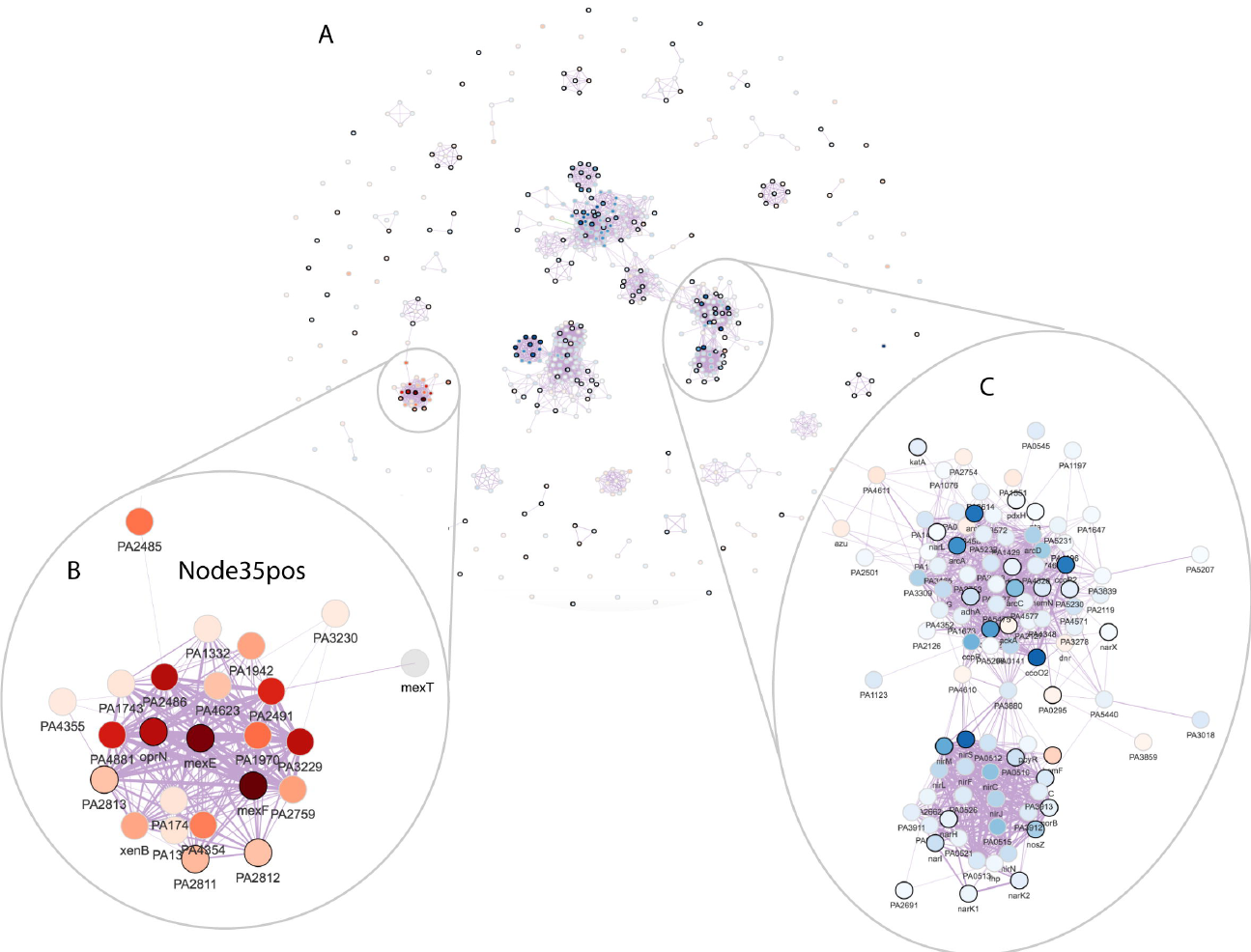
The gene-gene network for the example dataset. A: The ADAGE gene-gene network subset by genes in the selected active signatures. Each vertex in the network is a gene. Vertex color corresponds to gene expression fold change between *Δanr* mutant and wild-type (red: high in *Δanr* mutant; blue: low in *Δanr* mutant). Genes with black border are genes that have KEGG pathway annotations. The thickness of edges reflects how strong gene-gene relationships are in the ADAGE model. B: The network module of genes in Node35pos. C: The network module of genes involved in denitrification.

We next performed analyses designed to suggest the biological basis of the activated signatures. We linked signatures to KEGG pathways through enrichment tests. Twelve of the nineteen signatures were significantly enriched for one or more of fourteen KEGG pathways (Table 1, pathway association significance). We then tested whether these KEGG pathways were differentially active between wild-type and *Δanr* mutants using only genes shared by signatures and their associated pathways. Genes in seven of the fourteen KEGG pathways were significantly activated (adjusted p-value <=0.05) (Table 1, pathway activation significance). As a comparison, we also performed the popular gene set enrichment analysis (GSEA) [43] and considered pathways with FDR q-values lower than 0.05 in GSEA’s permutation test (Table 2). Five pathways were detected by both GSEA and ADAGE: Type VI secretion system; Cytochrome c oxidase, cbb3-type; Nitrogen metabolism; Denitrification, nitrate => nitrogen; Biosynthesis of siderophore group nonribosomal peptides. These pathways have been shown to be regulated by Anr [44]. Eight pathways were only detected by GSEA (Table 2, pathways with white background). Many are large pathways including the Ribosome and bacterial secretion system pathways (size of 56 and 90 respectively). GSEA has been found to be biased towards large gene sets [45]. Three of the GSEA-only pathways were associated with signatures that nearly met the signature selection criteria (Figure S2), and two were not associated with any ADAGE signatures. Nine KEGG pathways were not significantly enriched in GSEA but were associated with active ADAGE signatures (Table 1, pathways with white background). Among them, two pathways (Cyanoamino acid metabolism and Iron complex transport system) were considered activated by ADAGE signature analysis and also achieved high enrichment scores in GSEA (Table 2). The other seven pathways did not appear activated in this dataset. This primarily occurred when one signature was associated with several KEGG pathways, though sometimes only one or two of the pathways were strongly activated in this dataset. The fact that several pathways are grouped into one signature indicates that their expression frequently co-varies across the compendium. Therefore, pathways that are associated with active signatures but are not active themselves in a dataset should be viewed critically. They could be co-regulated pathways whose activities did not peak at the time of experiment or pathways that are co-regulated but only under conditions not relevant to this experiment.

**Table 1:** Signatures selected by ADAGE signature analysis for the example dataset and their associated KEGG pathways.

| Signature | Pathway | Pathway association significance (adj.p.val) | Pathway activation significance (adj.p.val) |
| --- | --- | --- | --- |
| Node276neg | KEGG-Module-M00156: Cytochrome c oxidase, cbb3-type | 1.10E-03 | 0.00E+00* |
| Node34pos | KEGG-Module-M00156: Cytochrome c oxidase, cbb3-type | 4.20E-03 | 1.00E-04* |
| Node38neg | KEGG-Module-M00334: Type VI secretion system | 1.00E-04 | 2.00E-04* |
| Node33neg | KEGG-Module-M00334: Type VI secretion system | 0.00E+00 | 2.00E-04* |
| Node28neg | KEGG-Pathway-pae00910: Nitrogen metabolism - Pseudomonas aeruginosa PAO1 | 1.00E-04 | 2.00E-04* |
| Node31pos | KEGG-Module-M00334: Type VI secretion system | 0.00E+00 | 2.00E-04* |
| Node28neg | KEGG-Module-M00529: Denitrification, nitrate => nitrogen | 1.00E-04 | 4.00E-04* |
| Node276neg | KEGG-Pathway-pae00460: Cyanoamino acid metabolism - Pseudomonas aeruginosa PAO1 | 2.10E-03 | 9.00E-04* |
| Node57neg | KEGG-Module-M00240: Iron complex transport system | 2.00E-04 | 7.50E-03* |
| Node155neg | KEGG-Module-M00240: Iron complex transport system | 4.50E-03 | 1.48E-02* |
| Node155neg | KEGG-Pathway-pae01053: Biosynthesis of siderophore group nonribosomal peptides - Pseudomonas aeruginosa PAO1 | 0.00E+00 | 1.50E-02* |
| Node57neg | KEGG-Pathway-pae01053: Biosynthesis of siderophore group nonribosomal peptides - Pseudomonas aeruginosa PAO1 | 0.00E+00 | 1.51E-02* |
| Node66pos | KEGG-Pathway-pae01053: Biosynthesis of siderophore group nonribosomal peptides - Pseudomonas aeruginosa PAO1 | 0.00E+00 | 1.52E-02* |
| Node205neg | KEGG-Pathway-pae01053: Biosynthesis of siderophore group nonribosomal peptides - Pseudomonas aeruginosa PAO1 | 0.00E+00 | 1.53E-02* |
| Node27neg | KEGG-Pathway-pae01053: Biosynthesis of siderophore group nonribosomal peptides - Pseudomonas aeruginosa PAO1 | 2.40E-03 | 1.59E-02* |
| Node33neg | KEGG-Module-M00644: Vanadium resistance, efflux pump MexGHI-OpmD | 3.00E-04 | 2.88E-01 |
| Node155neg | KEGG-Module-M00242: Zinc transport system | 1.35E-02 | 1.00E+00 |
| Node276neg | KEGG-Pathway-pae01503: Cationic antimicrobial peptide (CAMP) resistance - Pseudomonas aeruginosa PAO1 | 0.00E+00 | 1.00E+00 |
| Node276neg | KEGG-Pathway-pae00520: Amino sugar and nucleotide sugar metabolism - Pseudomonas aeruginosa PAO1 | 2.40E-03 | 1.00E+00 |
| Node276neg | KEGG-Module-M00721: Cationic antimicrobial peptide (CAMP) resistance, arnBCADTEF operon | 0.00E+00 | 1.00E+00 |
| Node9pos | KEGG-Pathway-pae00260: Glycine, serine and threonine metabolism - Pseudomonas aeruginosa PAO1 | 5.10E-03 | 1.00E+00 |
| Node9pos | KEGG-Pathway-pae00630: Glyoxylate and dicarboxylate metabolism - Pseudomonas aeruginosa PAO1 | 6.20E-03 | 1.00E+00 |

**Table 2:** GSEA result for the example dataset.

| Pathway | Pathway size | Enrichment score | Enrichment significance (FDR q-val) |
| --- | --- | --- | --- |
| KEGG-MODULE-M00178: RIBOSOME, BACTERIA | 56 | 0.622 | 0.00E+00* |
| KEGG-PATHWAY-PAE03010: RIBOSOME - PSEUDOMONAS AERUGINOSA PAO1 | 56 | 0.622 | 0.00E+00* |
| KEGG-MODULE-M00334: TYPE VI SECRETION SYSTEM | 43 | -0.83 | 0.00E+00* |
| KEGG-PATHWAY-PAE03070: BACTERIAL SECRETION SYSTEM - PSEUDOMONAS AERUGINOSA PAO1 | 90 | -0.609 | 4.29E-04* |
| KEGG-PATHWAY-PAE01053: BIOSYNTHESIS OF SIDEROPHORE GROUP NONRIBOSOMAL PEPTIDES - PSEUDOMONAS AERUGINOSA PAO1 | 6 | -0.995 | 1.13E-03* |
| KEGG-MODULE-M00331: TYPE II GENERAL SECRETION PATHWAY | 33 | -0.664 | 3.35E-03* |
| KEGG-PATHWAY-PAE00910: NITROGEN METABOLISM - PSEUDOMONAS AERUGINOSA PAO1 | 37 | -0.641 | 3.54E-03* |
| KEGG-MODULE-M00156: CYTOCHROME C OXIDASE, CBB3-TYPE | 8 | -0.877 | 7.24E-03* |
| KEGG-MODULE-M00332: TYPE III SECRETION SYSTEM | 18 | -0.722 | 9.69E-03* |
| KEGG-MODULE-M00529: DENITRIFICATION, NITRATE => NITROGEN | 10 | -0.81 | 1.06E-02* |
| KEGG-MODULE-M00023: TRYPTOPHAN BIOSYNTHESIS, CHORISMATE => TRYPTOPHAN | 9 | -0.808 | 2.99E-02* |
| KEGG-PATHWAY-PAE00280: VALINE, LEUCINE AND ISOLEUCINE DEGRADATION - PSEUDOMONAS AERUGINOSA PAO1 | 48 | -0.562 | 3.64E-02* |
| KEGG-PATHWAY-PAE00970: AMINOACYL-TRNA BIOSYNTHESIS - PSEUDOMONAS AERUGINOSA PAO1 | 27 | 0.623 | 4.38E-02* |
| KEGG-PATHWAY-PAE00460: CYANOAMINO ACID METABOLISM - PSEUDOMONAS AERUGINOSA PAO1 | 11 | -0.675 | 2.04E-01 |
| KEGG-MODULE-M00240: IRON COMPLEX TRANSPORT SYSTEM | 10 | -0.661 | 2.63E-01 |

ADAGE built not only signatures resembling existing pathways but also novel signatures that are unavailable in traditional pathway analysis. Seven differentially active signatures were not enriched for KEGG pathways, including Node35pos (Figure 5B), the most active signature in the *Δanr* mutant (Figure 4A). We also attempted to interpret Node35pos using GO terms but, as with our KEGG analysis, found it associated with no existing GO terms. Node35pos contains genes *mexEF* and *oprN*, which encode multidrug efflux protein, and many uncharacterized genes. The majority of the genes in Node35pos were highly expressed in *Δanr* mutants. To examine whether or not Node35pos captured a regulatory module, we analyzed the three most highly differentially expressed uncharacterized genes (PA4881, PA3229, and PA2486) in the STRING network [46] and found that they all returned similar networks that were subsets of Node35pos (Figure S3). Interestingly, these STRING networks were primarily built upon text mining. The reference papers used by STRING that performed a transcriptional profiling either did not release data [47] or used RNA-seq [48] and thus they were not included in the eADAGE model training set. Therefore, we concluded that signature Node35pos captured a transcriptional process, which was independently supported by the STRING network. MexEF-OprN is known to be regulated by MexT [49], which is connected to genes in Node35pos but is not in Node35pos itself (Figure 5B). Further examination showed that genes in Node35pos overlapped the MexT regulon (FDR q-value of 8.7e-23, MexT regulon obtained from CollecTF database [50]). The link between MexT and Anr has not been explicitly studied before. In strains lacking *anr*, the expression of *mexT* and MexT-regulated genes was higher. Because it has been shown that the MexT regulon, including the *mexEF-oprN* operon, is induced in response to nitrosative stress [51], we predict that the lack of the Anr-regulated denitrification genes *nar*, *nir*, and *nor* reduced detoxification of endogenously generated reactive nitrogen species [52–54], thereby activating MexT. The strong up-regulation of MexT, a redox-responsive regulator, may also be more active in *Δanr* mutant due to other changes in intracellular redox [55].

Through examining the overlapping genes in the seven uncharacterized signatures, we divided them into two groups (Figure S4). Group 1 contains MexT regulatory programs as represented by Node35pos. Group 2 contains many quorum sensing controlled genes, which are lower in the Δ*anr* mutant. Interestingly, the visualization of these pathways in output generated by this tool prompted the examination of connections between the MexT regulon and quorum sensing. Indeed, high expression of *mexEF-oprN* is associated with decreased quorum sensing due to the efflux of the QS molecule HHQ [56]. At the time we retrieved KEGG pathways and performed this analysis, signatures in Group 2 were still uncharacterized in KEGG. An updated analysis showed that signatures in Group 2 were now associated with KEGG pathways quorum sensing and phenazine biosynthesis, which were added to KEGG on 8-1-2016 and 3-27-2017 respectively. Signatures in Group 1 were still uncharacterized in the updated analysis. Quorum sensing and phenazine biosynthesis have been studied for a long time in *P. aeruginosa* with many well-characterized genes. The time lag in their annotation hinders their usage in traditional pathway analysis, yet ADAGE identified them directly from public data and grouped them into signatures. This again highlights the strength of ADAGE-based signature analysis: it does not rely on pre-defined pathways but uses regulatory patterns directly extracted from large compendia of gene expression data.

## Discussion

Researchers performing ADAGE signature analysis reverse the steps of traditional gene set analysis: they first identify signatures with statistically significant differential expression patterns before attempting interpretation. These researchers then only need to focus on signatures relevant to their experiments. This is important for organisms with incomplete or absent gene sets, because the alternative would be to curate every possible gene set before performing any analysis. The lag from discovery to gene set annotation also hinders the application of traditional gene set analysis to organisms with available curated resources. Our analysis revealed that even well characterized process, e.g. quorum sensing, can lack annotation in some resources and thus would not be detectable in traditional gene set analysis. However, biologists working in a field can often readily identify and interpret these differentially active ADAGE signatures. ADAGE signatures may also be comprised entirely of uncharacterized genes. Though such signatures would be difficult to interpret, they may represent novel biological processes. Thus ADAGE signature analysis is well suited to hypothesis generation in organisms about which little is known.

Constructing high-quality signatures in an unsupervised manner requires two key components: sufficient data and suitable feature extraction algorithms. The ideal data compendium should be a broad survey of an organism probed under many conditions. Signature analysis is unlikely to detect pathways that have never been perturbed in a compendium, and a heavily biased compendium would result in limited detection of biological processes. Though it is difficult to directly measure data comprehensiveness, both data quantity and a broad set of contributing research groups are expected to positively correlate with comprehensiveness. Quantity is important because more overall conditions are likely to have been measured, and the number of contributing research groups is important because they are likely to be studying different aspects of an organism’s biology. As genome-wide measurements continue to grow, we expect such methods to be more broadly applied to reveal perturbed biological processes and pathways.

Good feature extraction algorithms are also needed to best utilize the available data. Many feature extraction approaches have been applied on biological data, such as PCA [57–59], ICA [60–62], and NMF [63–65]. We previously developed ADAGE, a neural network-based approach, and found it to outperform PCA and ICA in representing biological states [9] and capturing KEGG pathways [12] in *P. aeruginosa*. However, it is still challenging to comprehensively evaluate the “effectiveness” of features built by different approaches, especially for less-studied organisms. Because every method has its own underlying assumptions and objectives, we expect them to learn different types of features and complement each other. The concept of gene set analysis with data-defined gene sets is not limited to ADAGE signatures. Future work will focus on expanding this analysis pipeline to more feature types and providing support for more organisms.

## Conclusions

Gene set analysis has been a powerful tool for interpreting the results of high throughput experiments. However, curating biologically important gene sets requires tremendous effort. As a result, many organisms, including those that are widely used to study certain processes, remain unannotated. In contrast with the sparsity of gene set annotations, high-throughput experimental data has accumulated rapidly. Unsupervised analysis of these data can reveal gene sets that parallel human-curated biological pathways. We introduced ADAGE signature analysis, a gene set analysis method powered by signatures built directly from expression data by ADAGE or other unsupervised feature construction methods. The approach includes three major steps: data preparation, differentially active signature detection, and signature interpretation. We compared this approach with GSEA on an example dataset and observed that ADAGE signature analysis and GSEA detected similar curated KEGG pathways. However, ADAGE signature analysis also identified a novel regulatory relationship unannotated in KEGG. This result highlights the advantage of ADAGE signature analysis: it does not depend on curated knowledgebases but instead the breadth of existing public data. ADAGE signature analysis is implemented in an R package and a web server for users with different backgrounds and needs. For those without a specific dataset to analyze, we also provided a gene-gene network view to explore transcriptional regulatory modules learned by ADAGE.

**List of abbreviations**
ADAGE: Analysis using Denoising Autoencoders for Gene Expression data
CFBE: Cystic Fibrosis genotype bronchial epithelial cells
GO: Gene Ontology
ICA: Independent Component Analysis
KEGG: Kyoto Encyclopedia of Genes and Genomes NMF: Non-negative Matrix Factorization
PCA: Principal Component Analysis
RMA: Robust Multi-array Average

## Declarations

**Ethics approval and consent to participate**

Not applicable

**Consent for publication**

Not applicable

**Availability of data and material**

The *anr* mutant dataset analyzed during the current study are available in the GEO repository, https://www.ncbi.nlm.nih.gov/geo/query/acc.cgi?acc=GSE67006.

## Authors’ contributions

JT and CSG conceived and designed the method. JT built the ADAGEpath R package and performed computational analyses. MH, DH, and RAZ developed the ADAGE web server. DAH analyzed the biological relevance of ADAGE signatures. JT, DAH, and CSG wrote the manuscript. All authors read and approved the final manuscript.

## Acknowledgements

The authors would like to thank Greg Way, Jaclyn Taroni, and Kurt Wheeler for helpful code review. The authors also would like to thank Georgia Doing for testing the R package.

---

Figure S1: The relationship between corruption level used in building ADAGE models and the redundancy of signatures derived from the models. The plot summarized results from 100 ADAGE models built at each corruption level.

A: The number of signature pairs with gene compositions significantly overlapped increases with corruption level until the corruption level reaches 30%.

B: As corruption level increases, the number of signatures in a model that enriched of the same KEGG pathway also increases on average, indicating the signatures become more redundant.

C: ADAGE models tend to capture more unique KEGG pathways (pathway coverage) when more noise was added during training until the corruption level is higher than 25%.

Figure S2: KEGG pathways enriched in the GSEA analysis but not associated with selected signatures in the ADAGE signature analysis. In the same volcano plot as Figure 4A, signatures associated with GSEA-only pathways are highlighted in blue while signatures lie on the first 10 Pareto fronts are highlighted in Red. Node137pos, Node252neg, and Node113neg obtained high significances in the activation test and would be considered if we lose the activation cutoff. Pathways Tryptophan biosynthesis, chorismate => tryptophan (KEGG-Module-M00023) and Aminoacyl-trna biosynthesis (KEGG-Pathway-pae00970) are not associated with any signature, so they were not labeled in the plot.

Figure S3: Validation of Node35pos as a transcriptional program via the STRING network. A: The largest connected module of the gene-gene network subset by genes in Node35pos. B: Gene-gene networks returned by STRING when searching PA2486, PA3229, and PA4881 respectively. The STRING networks of the three genes are subsets of the Node35pos network.

Figure S4: Groups of signatures that are uncharacterized by KEGG.

A: The signature similarity heatmap of uncharacterized signatures. Heatmap color reflects the odds ratio that two signatures overlap in their gene contents. Signatures are divided into two groups based on their similarity.

B: The largest connected module in the gene-gene network subset by genes in Group1 signatures. This module contains the MexT regulatory program.

C: The largest connected module in the gene-gene network subset by genes in Group2 signatures. This module contains many genes involved in quorum sensing.

